## Supplemental materials for "Dendrite intercalation between epidermal cells tunes nociceptor sensitivity to mechanical stimuli in *Drosophila* larvae"

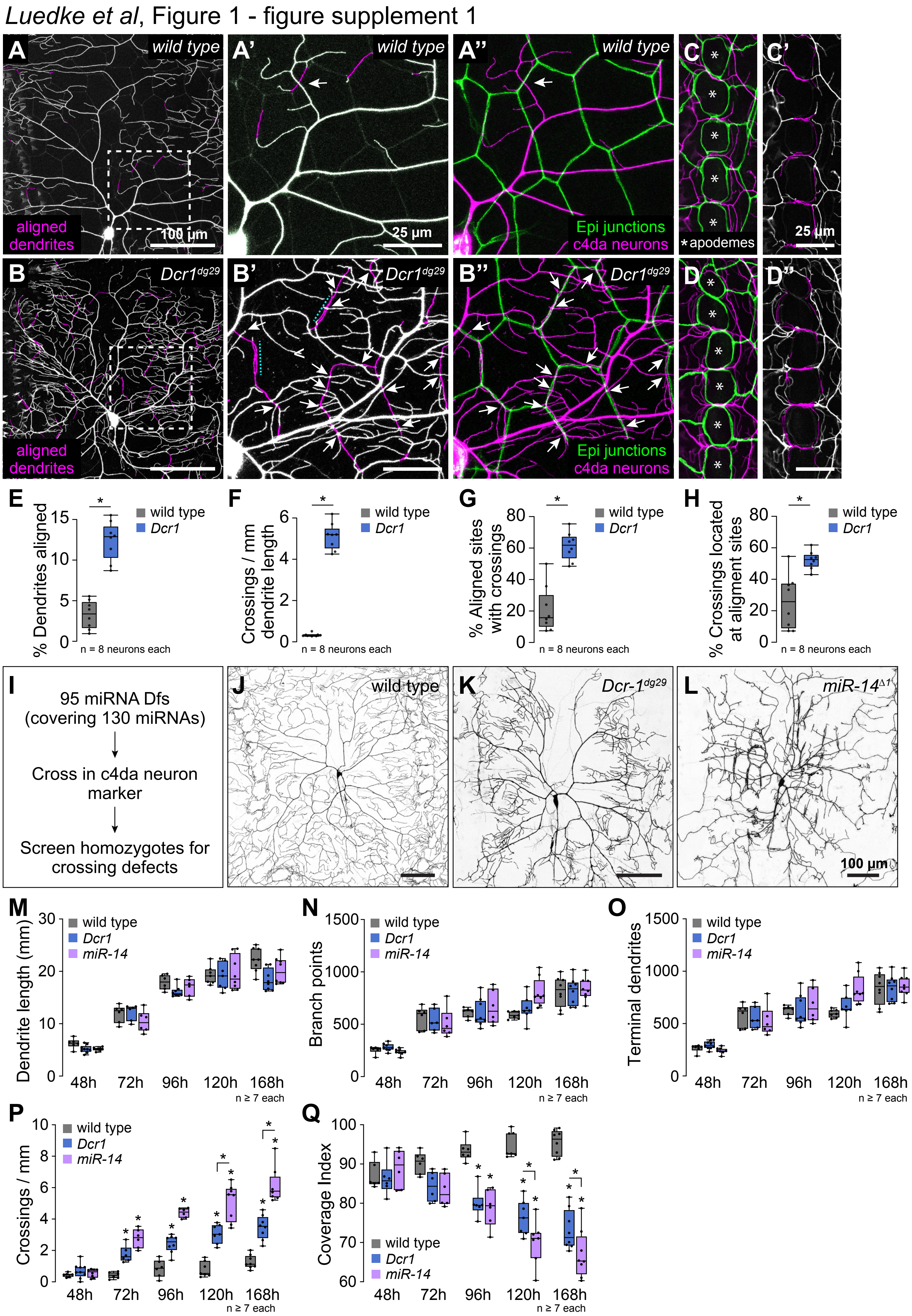

Luedke et al, Figure 1 - figure supplement 1
A
wild type
aligned dendrites
100 µm
A'
wild type
25 µm
A''
wild type
Epi junctions
c4da neurons
C
C'
*apodemes
25 µm
B
Dcr1dg29
aligned dendrites
B'
Dcr1dg29
B''
Dcr1dg29
Epi junctions
c4da neurons
D
D''
E
% Dendrites aligned
wild type
Dcr1
n = 8 neurons each
F
Crossings / mm dendrite length
wild type
Dcr1
n = 8 neurons each
G
% Aligned sites with crossings
wild type
Dcr1
n = 8 neurons each
H
% Crossings located at aligment sites
wild type
Dcr1
n = 8 neurons each
I
95 miRNA Dfs (covering 130 miRNAs)
Cross in c4da neuron marker
Screen homozygotes for crossing defects
J
wild type
K
Dcr-1dg29
L
miR-14Δ1
100 µm
M
Dendrite length (mm)
wild type
Dcr1
miR-14
48h 72h 96h 120h 168h
n ≥ 7 each
N
Branch points
wild type
Dcr1
miR-14
48h 72h 96h 120h 168h
n ≥ 7 each
O
Terminal dendrites
wild type
Dcr1
miR-14
48h 72h 96h 120h 168h
n ≥ 7 each
P
Crossings / mm
wild type
Dcr1
miR-14
48h 72h 96h 120h 168h
n ≥ 7 each
Q
Coverage Index
wild type
Dcr1
miR-14
48h 72h 96h 120h 168h
n ≥ 7 each

*Luedke et al*, Figure 1 - figure supplement 2

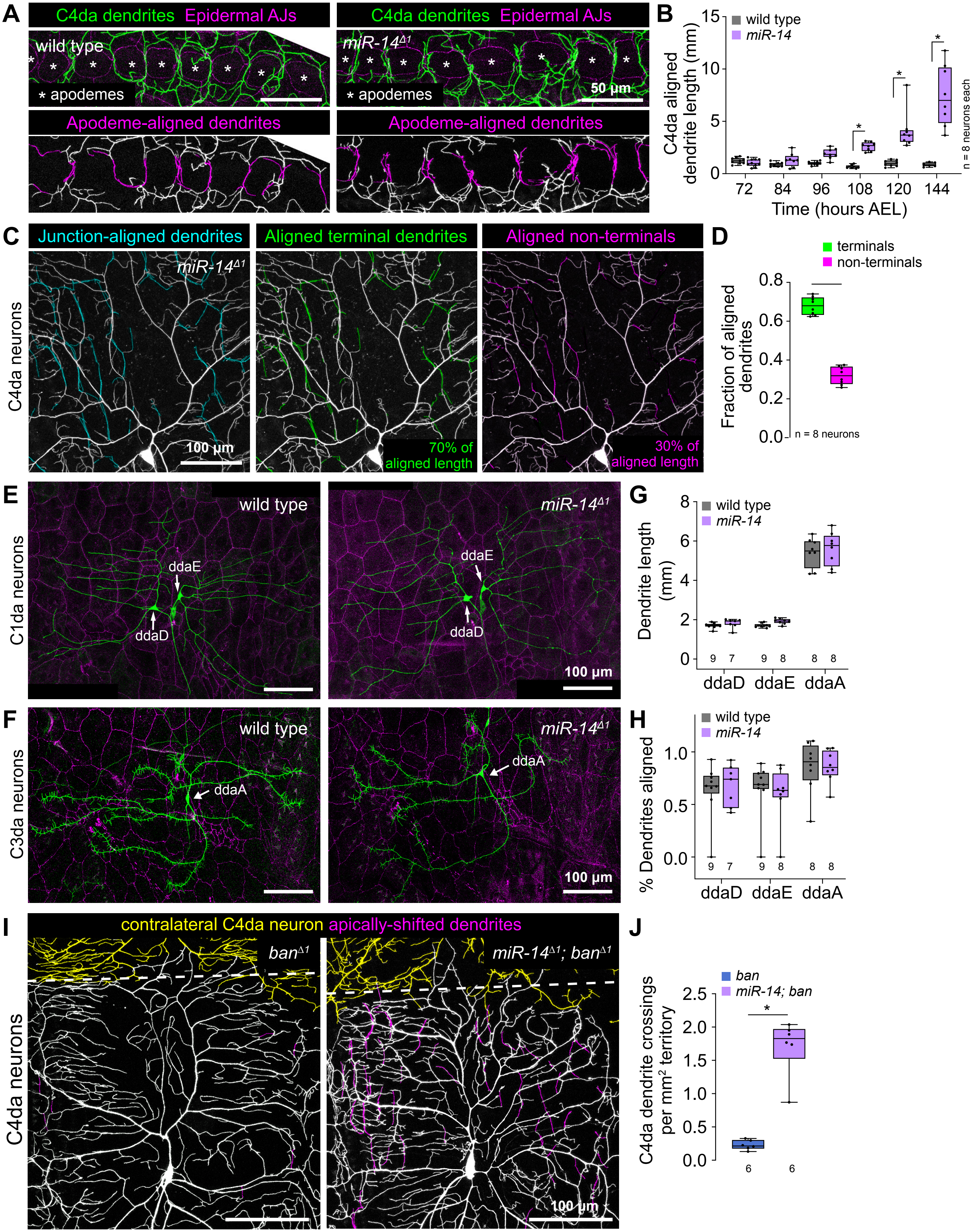

*Luedke et al*, Figure 2 - figure supplement 1

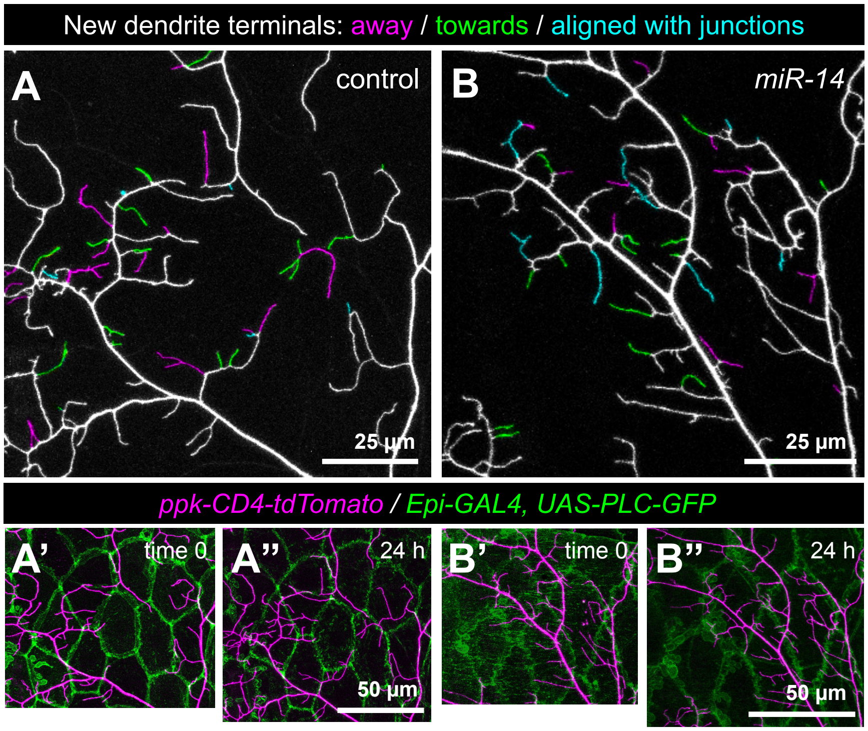

*Luedke et al*, Figure 4 - figure supplement 1

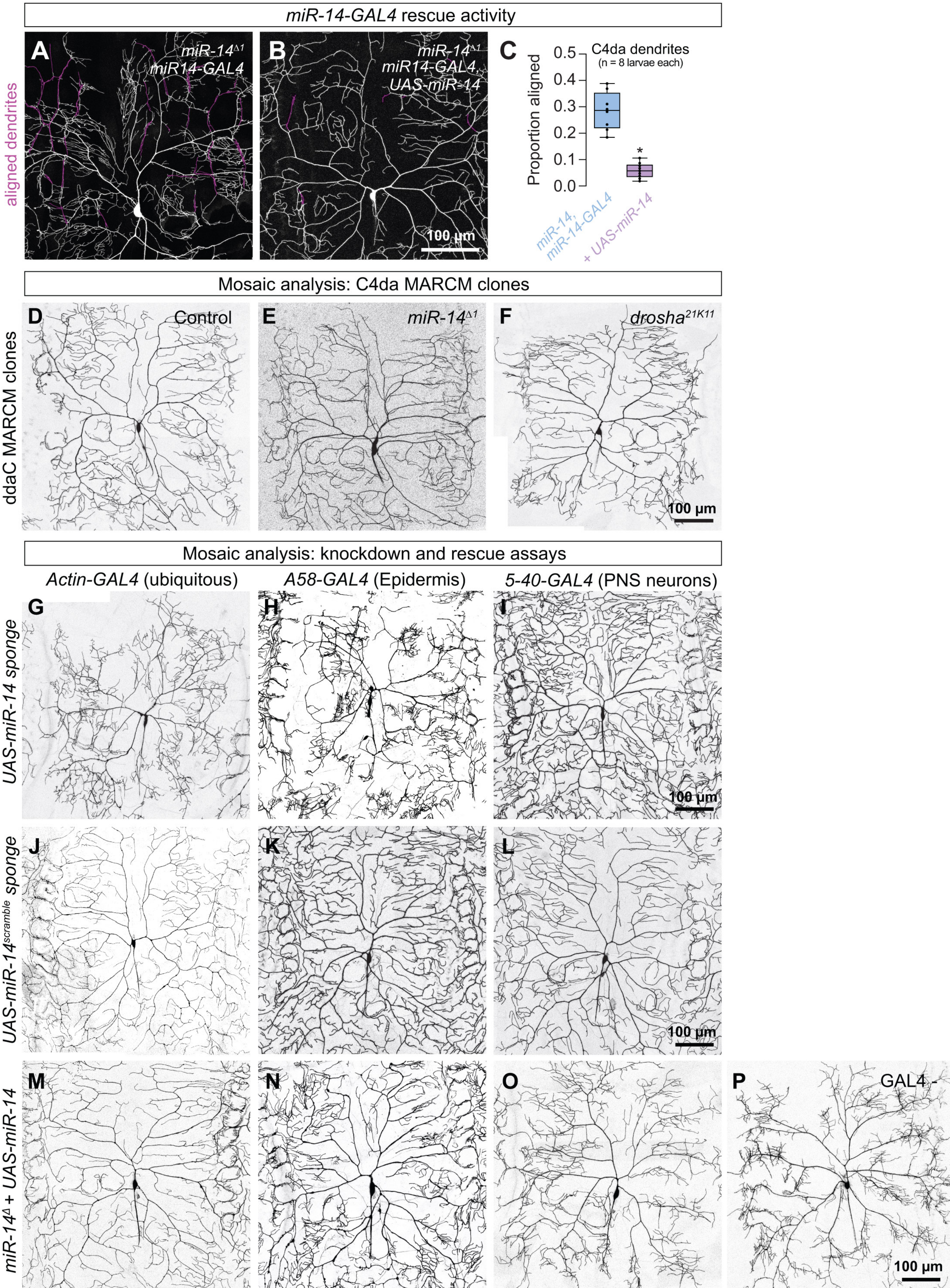

*Luedke et al*, Figure 5 - figure supplement 1

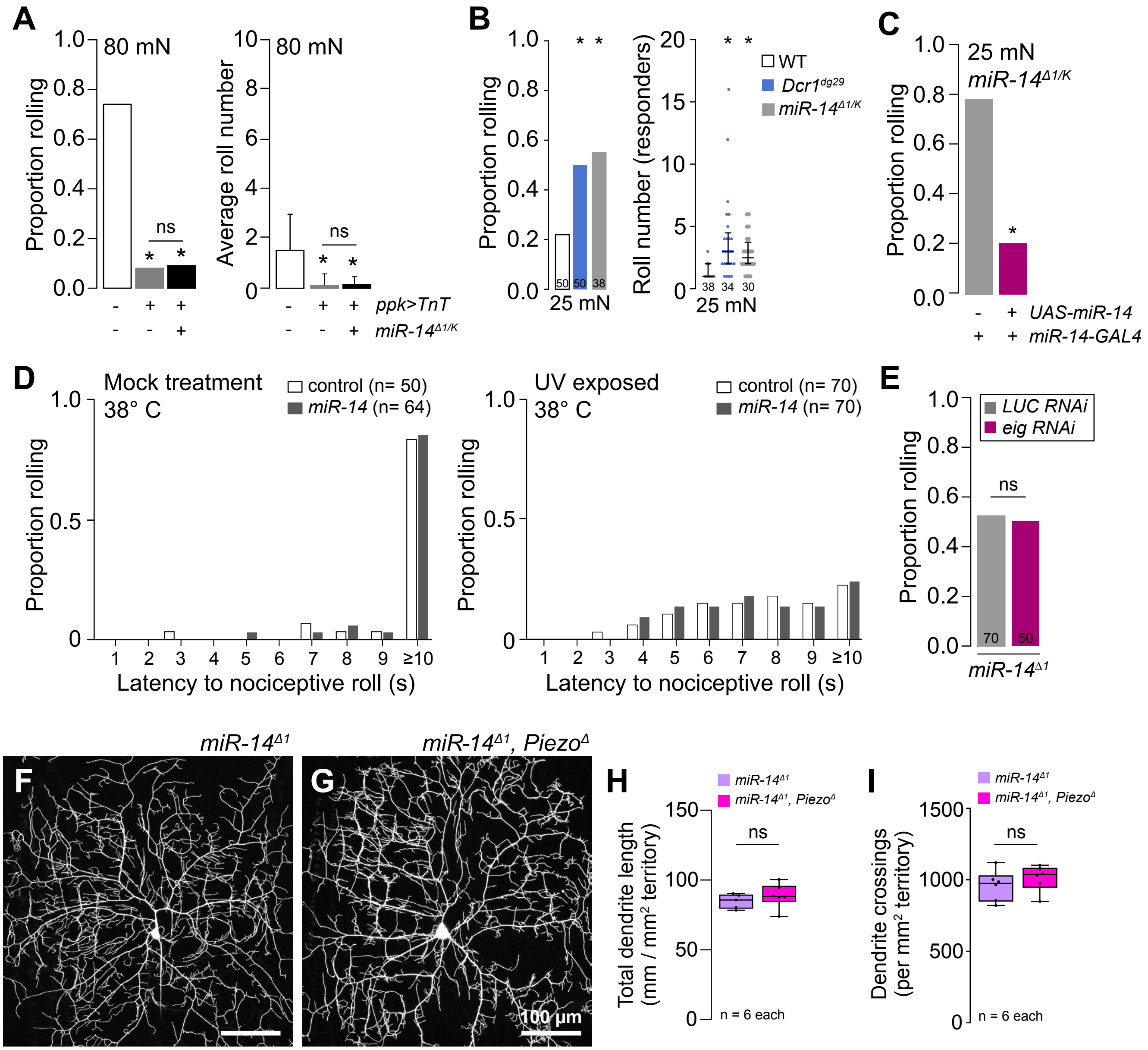

*Luedke et al*, Figure 6 - figure supplement 1

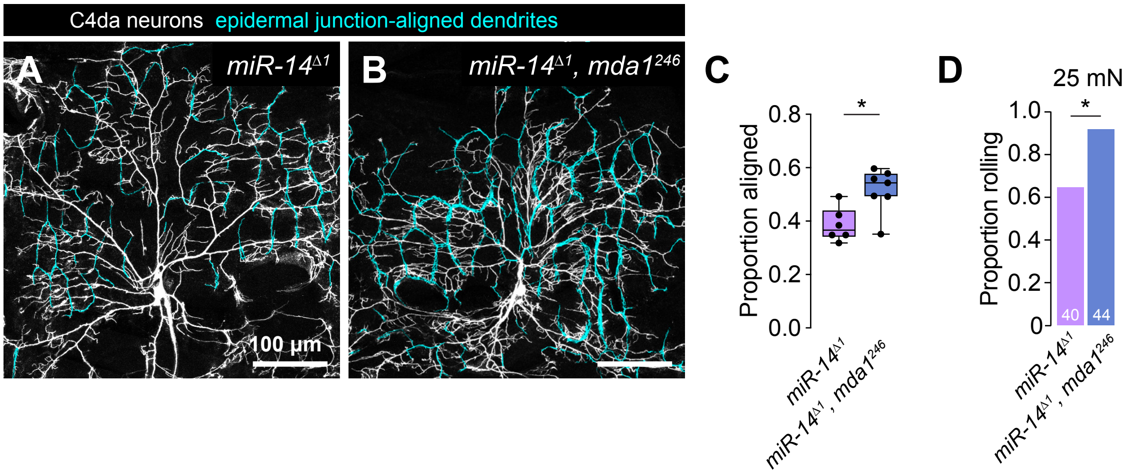

*Luedke et al*, Figure 7 - figure supplement 1

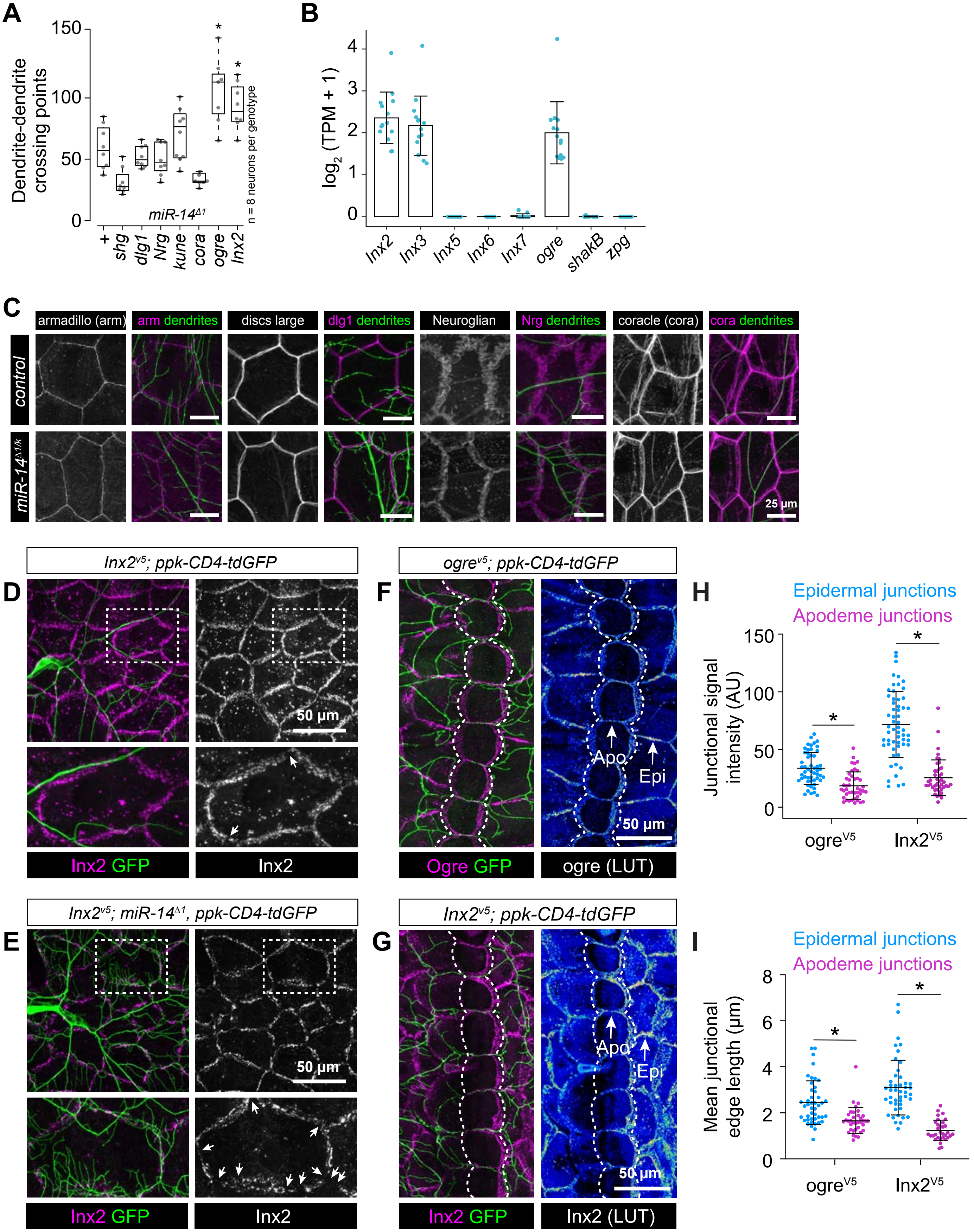

*Luedke et al,* Figure 7 - figure supplement 2

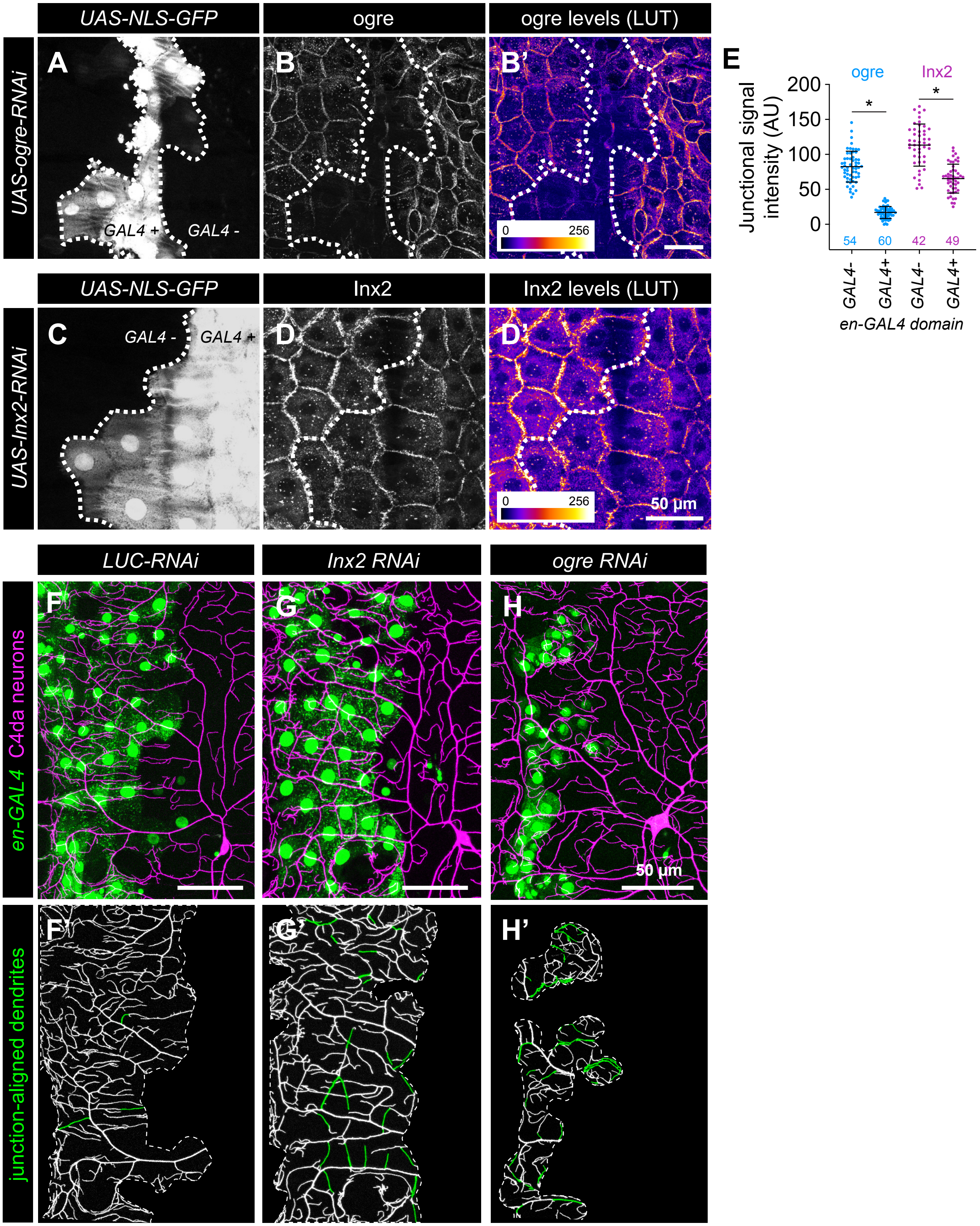

**Table S1. Alleles used in this study**

| Allele | Uses / Features | Identifier (RRID) |
|---|---|---|
| *5-40-GAL4* | GAL4 driver (PNS neurons) | Flybase_FBal0221791 |
| *98b-GAL4* | GAL4 driver (C1da neurons) | Flybase_FBal0305321 |
| *Act5C-GAL4* | GAL4 driver (ubiquitous) | BDSC_3954 |
| *A58-GAL4* | GAL4 driver (epidermis) | Flybase_FBal0181674 |
| *en2.4-GAL4* | GAL4 driver (epidermis) | BDSC_30564 |
| *miR-14-GAL4* | GAL4 driver (*miR-14* expression domain) | This study |
| *NompC-GAL4* | GAL4 driver (C3da neurons) | BDSC_36361 |
| *ppk-GAL4* | GAL4 driver (C4da neurons) | BDSC_32079 |
| *R38F11-GAL4* | GAL4 driver (epidermis) | BDSC_50014 |
| *sr-GAL4* | GAL4 driver (apodemes) | BDSC_26663 |
| *ppk-LexA* | LexA driver (C4da neurons) | Flybase_FBal0336342 |
| *UAS-mCherry.scramble.sponge* | Knockdown (control miRNA sponge) | BDSC_61501 |
| *UAS-mCherry.miR-14.sponge* | Knockdown (*miR-14* sponge) | BDSC_61382 |
| $UAS\text{-}eiger^{IR}$ | Knockdown (*eiger* RNAi line) | BDSC_58993 |
| *UAS-Inx2RNAi* | Knockdown (*Inx2* RNAi line) | BDSC_42645 |
| *UAS-LUC* | Knockdown (*Luciferase* RNAi line) | BDSC_31603 |
| *UAS-ogreRNAi* | Knockdown (*ogre* RNAi line) | BDSC_44048 |
| *FRT42D* | MARCM reagent | BDSC_1802 |
| *SOP-FLP* | MARCM reagent | Flybase_FBal0278148 |
| *Tub-GAL80* | MARCM reagent | BDSC_9917 |
| $ban^{\Delta 1}$ | Mutant allele | BDSC_58878 |
| $cora^{5}$ | Mutant allele | BDSC_52233 |
| $Dcr1^{mn29}$ | Mutant allele | This study |
| $dlg1^{A}$ | Mutant allele | BDSC_57086 |
| $Drosha^{21K11}$ | Mutant allele | Flybase_FBal0268190 |
| $inx2^{G0118}$ | Mutant allele | BDSC_11826 |
| $kune^{C309}$ | Mutant allele | BDSC_16333 |
| $mdi1^{242}$ | Mutant allele | This study |
| $mdi2^{246}$ | Mutant allele | This study |
| $mdi3^{442}$ | Mutant allele | This study |
| $mdi4^{51}$ | Mutant allele | This study |
| $miR\text{-}14^{\Delta 1}$ | Mutant allele | BDSC_33067 |
| $miR\text{-}14^{k10213}$ | Mutant allele | BDSC_10982 |
| $nan^{GAL4}$ | Mutant allele | BDSC_68205 |
| $NompC^{1}$ | Mutant allele | BDSC_42260 |
| $NompC^{3}$ | Mutant allele | BDSC_42258 |
| $Nrg^{14}$ | Mutant allele | BDSC_5708 |
| $ogre^{1}$ | Mutant allele | Flybase_FBal0013231 |
| $pain^{1}$ | Mutant allele | BDSC_27895 |
| $piezo^{KO}$ | Mutant allele | BDSC_58770 |
| $ppk^{ESB}$ | Mutant allele | BDSC_79622 |
| $shg^{2}$ | Mutant allele | BDSC_3085 |
| $Tig^{A1}$ | Mutant allele | BDSC_8795 |
| $Tig^{x}$ | Mutant allele | BDSC_8796 |
| $TrpA1^{1}$ | Mutant allele | BDSC_26263 |
| *UAS-if* | Overexpression/rescue construct | Flybase_FBal0062798 |
| *UAS-Inx2.Sb* | Overexpression/rescue construct | Flybase_FBal0338197 |
| *UAS-LUC-miR-14.T* | Overexpression/rescue construct | BDSC_41178 |
| *UAS-mew* | Overexpression/rescue construct | Flybase_FBal0062567 |
| *UAS-ogre.S* | Overexpression/rescue construct | Flybase_FBal0338192 |
| *UAS-TNT* | Overexpression/rescue construct | BDSC_28997 |
| $inx2^{V5}$ | Reporter (V5-tagged Inx2) | Flybase_FBal0345156 |
| $ogre^{V5}$ | Reporter (V5-tagged Ogre) | Flybase_FBal0345155 |
| *Nrg167GFP* | Reporter (epidermal septate junctions) | BDSC_6844 |

| | | |
|---|---|---|
| *Nrx-IV-GFP* | Reporter (epidermal septate junctions) | BDSC_50798 |
| *ppk-CD4-tdGFP$^{1b}$* | Reporter (C4da neurons) | BDSC_35842 |
| *ppk-CD4-tdTomato$^{10A}$* | Reporter (C4da neurons) | BDSC_35845 |
| *shg$^{mCherry}$* | Reporter (epidermal adherens junctions) | BDSC_59014 |
| *trol-GFP* | Reporter (epidermal BM) | Flybase_FBal0243610 |
| *Tub-GFP* | Reporter (control miRNA sensor) | FBtp0017439 |
| *Tub-GFP.mir-14* | Reporter (*miR-14* sensor) | FBtp0056974 |
| *UAS-CD4-tdGFP* | Reporter (membrane-targeted GFP) | BDSC_35839 |
| *UAS-GCaMP6s* | Reporter (calcium imaging) | BDSC_42749 |
| *AOP-GCaMP6s* | Reporter (calcium imaging) | BDSC_44273 |
| *UAS-Gerry* | Reporter (ratiometric calcium imaging) | BDSC_80141 |
| *UAS-mCD8-GFP* | Reporter (membrane-targeted GFP) | BDSC_5137 |
| *UAS-NLS-GFP* | Reporter (nuclear-targeted GFP) | BDSC_4776 |
| *UAS-PLC$^{\delta}$-PH-GFP* | Reporter (PIP2 reporter) | BDSC_39693 |
| *UAS-RedStinger$^{4}$* | Reporter (NLS-RFP) | BDSC_8546 |
| *UAS-tdTomato* | Reporter (RFP) | BDSC_36328 |
